# Behavioral variation across multiple phases of intravenous cocaine self-administration among genetically diverse mouse populations

**DOI:** 10.1101/2025.04.23.650259

**Authors:** Price E. Dickson, Udita Datta, Troy D. Wilcox, Ashley A. Auth, Robyn L. Ball, Matt Dunn, Heidi S. Fisher, Alyssa Klein, Michael R. Leonardo, Tyler A. Roy, Michael C. Saul, Jason A. Bubier, Leona H. Gagnon, Vivek M. Philip, Lisa M. Tarantino, James D. Jentsch, Elissa J. Chesler

## Abstract

Genetic and other predisposing factors can influence the progression from initiation of drug intake to compulsive substance use through distinct biobehavioral processes. Operant cocaine self-administration studies in laboratory mice offer a powerful method to dissect the biology of this progression from initiation, dose-response, extinction, and cued reinstatement in a controlled, tractable system. However, many such studies encompass limited genetic diversity and rarely examine self-administration behaviors beyond the acquisition stage. Here, we study three high-diversity mouse populations – 50 strains from the Collaborative Cross (CC) reference panel, a large sample of Diversity Outbred (J:DO) population and their eight founder strains – to characterize the varied phenotypic manifestation of behaviors across multiple phases of cocaine intravenous self-administration (IVSA) in both sexes. We observed distinct strain differences among the founders and CC strains in all phases of self-administration, with heritability estimates ranging from 0 to 0.585 and many CC and J:DO phenotypic values exceeding the range of founders including the C57BL/6J strain. Sex differences were common across behaviors, some manifesting as main effects, others as strain interactions. Finally, by adopting a multi-stage design, we identified extreme strains for various cocaine intake and response traits and evaluated whether these strains exhibited differences in behavioral assays that model compulsive drug seeking. Together, these findings demonstrate the utility of extended self-administration protocols in advanced mouse populations for discovery and characterization of biological mechanisms of substance use traits and for preclinical studies in relevant, complex mouse models.

## 1. Introduction

Cocaine use disorder (CUD) is a heritable disorder for which the underlying biological mechanisms remain largely unknown (Pierce et al., 2018); current treatment options are limited, and no FDA-approved pharmacotherapies are available (Buchholz & Saxon, 2019). Among the 1.9% of the US population who used cocaine in 2022, 0.5% of adults and 0.8% of young adults developed CUD (SAMHSA, 2023). Vulnerability to substance use disorders, including CUD, is associated with genetics, sex differences, and environmental factors and manifests in multiple ways including enhanced likelihood to initiate drug use, increased intake, persistent drug seeking, and propensity to relapse (Egervari et al., 2018; Swendsen & Le Moal, 2011).

Rodent studies of operant intravenous self-administration (IVSA) procedures have revealed biological mechanisms of behaviors relevant to human CUD (Huggett et al., 2021; Mews et al., 2023; Peltz & Tan, 2021) and this approach has high face validity (Sanchis-Segura & Spanagel, 2006; Spanagel, 2017; Vengeliene et al., 2008). By allowing animals to regulate their drug intake, IVSA supports longitudinal studies that model aspects of the substance use trajectory including initiation, maintenance of response across different drug doses, extinction of drug-associated behaviors when access to the drug is removed, and reinstatement with stress, priming injections, or drug-paired cues (Bossert et al., 2013; Dickson et al., 2011; Slosky et al., 2022). Most genetic studies of drug self-administration in mice focus primarily on acquisition, the earliest phase of drug taking (e.g., Bagley et al., 2022; Belin-Rauscent et al., 2016; Khan et al., 2023). However, by examining multiple phases in the same animal or within isogenic strains, researchers can identify biological mechanisms that vary through critical behavioral transitions, as shown in a survey of male mice from eight strains (Roberts et al., 2018). The vast majority of mouse IVSA studies have emphasized a few widely used strains and substrains, typically C57BL/6 and strains with mutations engineered on this background (Grahame et al., 1995; Thomsen & Caine, 2007; Yoshiki et al., 2022). The nature and extent of strain and sex differences across phases in larger, higher diversity populations remains unknown. IVSA studies in high diversity genetic reference populations with a broad range of behavioral diversity enable the integration of data across populations and species for discovery of the biological mechanisms underlying such variation (Ball et al., 2024; Reynolds et al., 2021).

Genetically diverse inbred strains, recombinant inbred lines, and outbred populations have been developed to identify causative genes and pathways underlying complex disease (Saul et al., 2019). The Collaborative Cross (CC) recombinant inbred panel (Churchill et al., 2004) and the Diversity Outbred (J:DO) population (Churchill et al., 2012) were derived from eight founder stains, including five classical laboratory strains (C57BL/6J, A/J, 129S1/SvImJ, NOD/ShiLtJ, and NZO/HILtJ) and three wild-derived strains (CAST/EiJ, PWK/PhJ, and WSB/EiJ), capturing nearly 90% of known genetic variation among mouse strains. The CC panel provides extensive phenotypic and genetic variation due to recombination of the founder genomes, resulting from the well-randomized breeding scheme employed to produce this population (Chesler et al., 2008). The inbred nature of the CC strains enables simplified integration of data across experiments, phenotypes, experimental conditions, and time (Churchill et al., 2004). Additionally, the strains in the CC mouse panel harbor variants across the genome, which when combined, produce a wider range of more vulnerable or resistant states of disease (Saul et al., 2019; Schoenrock et al., 2020).

The J:DO outbred population has segregating genetic variation derived from the same founder strains, making them highly heterozygous and providing genetic variation (∼87.5% of the genome) in virtually all genes in the genome. Continuous outbreeding ensures that each J:DO mouse is genetically unique. The ever-expanding number of genetic recombinations in the J:DO enables high precision quantitative trait locus (QTL) mapping (Dickson, Ndukum, et al., 2015; Logan et al., 2013). Collectively, these genetically diverse recombinant mouse strains and their founders are tractable models expressing a broad range of phenotypic variation.

In this study, we examined the breadth of multiple CUD-relevant IVSA phenotypes in male and female mice from CC and J:DO populations and their eight inbred founder strains. We assessed phenotypic variation and covariation in behaviors extending well beyond the initial phases of drug self-administration in (1) initiation of drug use behavior during acquisition of cocaine self-administration, (2) maintenance of drug seeking behaviors over multiple doses, (3) persistence in drug seeking behavior during an extinction phase when both drug and drug-paired cues were unavailable, and (4) the propensity to resume drug seeking in the presence of cues previously paired with drug taking during a cued reinstatement phase. From these observations, we defined the contribution of genetic factors for each trait through heritability estimates, evaluated the relationships among traits to reveal shared and distinct genetic mechanisms of trait regulation, and identified extreme strains that exhibit a strong propensity or resistance toward cocaine self-administration behaviors. Finally, we evaluated the impact of these differences on assays designed to model compulsive drug seeking.

## 2. Materials and Methods

### 2.1. Subjects

Mice tested for cocaine IVSA were adult (9 - 24 weeks old) from the eight founder strains of the CC/J:DO populations (N_female_ = 105, N_male_ = 90), 50 of the CC strains (N_female_ = 181, N_male_ = 201), and the J:DO population (N_female_ = 30, N_male_ = 35; Table 1). Subjects in the choice procedure were adults from two strains: CC002/UncJ (N_female_ = 8, N_male_ = 10) and CC003/UncJ (N_female_ = 7, N_male_ = 6), and in the punishment procedure, adults from three strains CC002/UncJ (N_female_ = 3, N_male_ = 5), CC003/UncJ (N_female_ = 2, N_male_ = 3), and NZO/HILtJ (N_female_ = 6, N_male_ = 6). All mice were obtained from Genetic Resource Science Repository (founder and CC mice) or JAX Mice Clinical and Research Services (J:DO mice) Production Colonies at The Jackson Laboratory, Bar Harbor, ME and transferred to the Research Animal Facility by wheeled cart. Mice were housed in same sex groups of three animals in duplex polycarbonate cages on ventilated racks which provided 99.997% HEPA filtered air to each cage (Maxi-Miser Interchangeable System, Thoren). Cage lids had filtered tops. Mice were maintained in a climate-controlled room under a 12:12 light-dark cycle (lights on at 0600 h). All testing occurred during the light phase. Bedding was changed weekly, and mice were provided free access to food (NIH31 5K52 chow, LabDiet/PMI Nutrition, St. Louis, MO) and acidified water. A nestlet and a Shepherd Shack were provided in each cage for enrichment. Prior to the IVSA procedure, mice were assessed on a novelty-related test battery (open field, light/dark, hole-board, and novelty place preference) described in (Dickson, Ndukum, et al., 2015). All procedures were approved by JAX Animal Care and Use Committee (AUS# 100007, Chesler, PI) and conducted in compliance with National Institutes of Health Guidelines for the Care and Use of Laboratory Animals 8^th^ Edition.

**Table 1.** Sample size by strain and sex.

| CC strains | JAX Stock ID# | MGI ID# | Total Mice Tested |  | Reached acquisition criteria |  | Completed IVSA testing |  | Positive Extinction Rate |  |
| --- | --- | --- | --- | --- | --- | --- | --- | --- | --- | --- |
|  |  |  | Female | Male | Female | Male | Female | Male | Female | Male |
| CC001/UncJ | 021238 | 5649079 | 4 | 4 | 3 | 4 | 3 | 4 | 2 | 3 |
| CC002/UncJ | 021236 | 5649080 | 3 | 2 | 3 | 2 | 3 | 2 | 1 | 2 |
| CC003/UncJ | 021237 | 5649081 | 4 | 4 | 1 | 3 | 2 | 4 | 1 | 4 |
| CC004/TauUncJ | 020944 | 5649082 | 12 | 22 | 12 | 21 | 9 | 19 | 4 | 14 |
| CC005/TauUncJ | 020945 | 5649083 | 5 | 3 | 4 | 3 | 4 | 2 | 4 | 1 |
| CC006/TauUncJ | 022869 | 5649237 | 3 | 2 | 2 | 1 | 2 | 1 | 2 | 0 |
| CC007/UncJ | 029625 | 5805789 | 3 | 4 | 3 | 3 | 3 | 3 | 2 | 2 |
| CC008/GeniUncJ | 026971 | 5659476 | 4 | 2 | 1 | 2 | 1 | 2 | 1 | 2 |
| CC009/UncJ | 018856 | 5649238 | 3 | 3 | 2 | 2 | 2 | 2 | 2 | 1 |
| CC010/GeniUncJ | 021889 | 5649239 | 1 | 2 | 0 | 0 | 0 | 0 | 0 | 0 |
| CC011/UncJ | 018854 | 5649240 | 6 | 5 | 4 | 5 | 2 | 3 | 2 | 2 |
| CC012/GeniUncJ | 028409 | 5694080 | 6 | 7 | 2 | 1 | 1 | 1 | 1 | 0 |
| CC013/GeniUncJ | 021892 | 5649241 | 3 | 3 | 3 | 3 | 3 | 3 | 1 | 2 |
| CC015/UncJ | 018859 | 5659478 | 3 | 3 | 2 | 2 | 2 | 2 | 2 | 2 |
| CC016/GeniUncJ | 024684 | 5659479 | 3 | 5 | 2 | 3 | 2 | 3 | 1 | 2 |
| CC017/UncJ | 022870 | 5649242 | 4 | 5 | 3 | 2 | 3 | 2 | 2 | 1 |
| CC018/UncJ | 021890 | 5649243 | 3 | 5 | 2 | 5 | 2 | 5 | 1 | 2 |
| CC019/TauUncJ | 021894 | 5649244 | 5 | 3 | 5 | 3 | 4 | 3 | 1 | 1 |
| CC020/GeniUncJ | 025129 | 5659480 | 2 | 2 | 2 | 1 | 2 | 1 | 2 | 1 |
| CC023/GeniUncJ | 025131 | 5659483 | 4 | 3 | 2 | 3 | 1 | 3 | 1 | 3 |
| CC024/GeniUncJ | 021891 | 5649245 | 3 | 3 | 1 | 1 | 1 | 1 | 1 | 0 |
| CC025/GeniUncJ | 018857 | 5649246 | 2 | 2 | 2 | 1 | 2 | 1 | 1 | 1 |
| CC026/GeniUncJ | 024685 | 5649247 | 6 | 4 | 5 | 3 | 5 | 3 | 5 | 3 |
| CC027/GeniUncJ | 025130 | 5659484 | 5 | 5 | 3 | 5 | 3 | 4 | 3 | 4 |
| CC028/GeniUncJ | 025126 | 5659485 | 5 | 6 | 5 | 6 | 4 | 5 | 2 | 4 |
| CC029/UncJ | 026972 | 5659486 | 2 | 2 | 1 | 2 | 0 | 2 | 0 | 2 |
| CC030/GeniUncJ | 025426 | 5659487 | 2 | 2 | 2 | 2 | 2 | 2 | 2 | 1 |
| CC032/GeniUncJ | 020946 | 5649248 | 3 | 4 | 3 | 4 | 3 | 4 | 3 | 3 |
| CC033/GeniUncJ | 025910 | 5659488 | 3 | 2 | 2 | 2 | 1 | 2 | 1 | 1 |
| CC035/UncJ | 024073 | 5659489 | 4 | 2 | 3 | 2 | 3 | 2 | 2 | 2 |
| CC036/UncJ | 025127 | 5659490 | 3 | 3 | 3 | 2 | 3 | 2 | 3 | 2 |
| CC037/TauUncJ | 025423 | 5649249 | 4 | 4 | 0 | 0 | 0 | 0 | 0 | 0 |
| CC040/TauUncJ | 023831 | 5649250 | 4 | 3 | 2 | 2 | 2 | 2 | 1 | 1 |
| CC041/TauUncJ | 021893 | 5649251 | 12 | 25 | 0 | 1 | 1 | 1 | 0 | 1 |
| CC043/GeniUncJ | 023828 | 5649256 | 2 | 3 | 2 | 3 | 2 | 1 | 1 | 1 |
| CC044/UncJ | 026426 | 5659493 | 3 | 3 | 0 | 0 | 0 | 0 | 0 | 0 |
| CC045/GeniUncJ | 025425 | 5659494 | 2 | 2 | 1 | 0 | 1 | 0 | 1 | 0 |
| CC051/TauUncJ | 021897 | 5649257 | 2 | 3 | 2 | 3 | 2 | 3 | 2 | 2 |
| CC057/UncJ | 024683 | 5659646 | 4 | 4 | 4 | 2 | 4 | 2 | 3 | 2 |
| CC059/TauUncJ | 025125 | 5659650 | 4 | 5 | 4 | 5 | 4 | 4 | 1 | 3 |
| CC060/UncJ | 026427 | 5659652 | 0 | 1 | 0 | 1 | 0 | 1 | 0 | 1 |
| CC061/GeniUncJ | 023826 | 5649258 | 2 | 2 | 2 | 2 | 2 | 2 | 2 | 2 |
| CC068/TauUncJ | 025908 | 5649259 | 5 | 3 | 1 | 0 | 0 | 0 | 0 | 0 |
| CC074/UncJ | 018855 | 5659666 | 3 | 4 | 1 | 2 | 1 | 2 | 0 | 2 |
| CC075/UncJ | 027293 | 5659668 | 4 | 3 | 1 | 1 | 1 | 1 | 1 | 1 |
| CC078/TauUncJ | 025989 | 5796475 | 2 | 3 | 2 | 2 | 2 | 3 | 1 | 2 |
| CC079/TauUncJ | 025990 | 5796476 | 2 | 2 | 2 | 2 | 2 | 2 | 1 | 2 |
| CC080/TauUncJ | 025988 | 5796471 | 3 | 2 | 2 | 1 | 2 | 2 | 2 | 1 |
| CC083/UncJ | 031921 | 6144025 | 2 | 1 | 1 | 1 | 1 | 1 | 0 | 1 |
| CC084/TauJ | 028923 | 6159255 | 2 | 4 | 2 | 3 | 2 | 3 | 2 | 2 |
| Total |  |  | 181 | 201 | 117 | 130 | 107 | 123 | 74 | 92 |

| Founder Strain | JAX Stock ID# | MGI ID# | Total Mice Tested |  | Reached acquisition criteria |  | Completed IVSA testing |  | Positive Extinction Rate |  |
| --- | --- | --- | --- | --- | --- | --- | --- | --- | --- | --- |
|  |  |  | Female | Male | Female | Male | Female | Male | Female | Male |
| A/J | 000646 | 2159747 | 12 | 9 | 9 | 9 | 7 | 7 | 7 | 7 |
| C57BL/6J | 000664 | 3028467 | 21 | 19 | 11 | 11 | 10 | 11 | 8 | 11 |
| 129S1/SvImJ | 002448 | 3037980 | 12 | 12 | 0 | 0 | 0 | 0 | 0 | 0 |
| NOD/ShiLtJ | 001976 | 2162056 | 14 | 10 | 12 | 9 | 10 | 6 | 9 | 5 |
| NZO/HiLtJ | 002105 | 2173835 | 14 | 11 | 13 | 11 | 10 | 10 | 9 | 8 |
| CAST/EiJ | 000928 | 2159793 | 8 | 4 | 8 | 4 | 1 | 2 | 1 | 2 |
| PWK/PhJ | 003715 | 2160654 | 10 | 11 | 10 | 11 | 6 | 6 | 6 | 6 |
| WSB/EiJ | 001145 | 2160667 | 14 | 14 | 2 | 7 | 1 | 5 | 1 | 1 |
| Total |  |  | 105 | 90 | 65 | 62 | 45 | 47 | 41 | 40 |

| Diversity Outbred | JAX Stock ID# | MGI ID# | Total Mice Tested |  | Reached acquisition criteria |  | Completed IVSA testing |  | Positive Extinction Rate |  |
| --- | --- | --- | --- | --- | --- | --- | --- | --- | --- | --- |
|  |  |  | Female | Male | Female | Male | Female | Male | Female | Male |
| J:DO | 009376 | 4412282 | 30 | 35 | 21 | 25 | 11 | 16 | 8 | 15 |

For IVSA drug delivery, mice were cannulated with a silicone catheter (CNC-2/3S-082109E/12, Access Technologies, Skokie, Illinois) implanted into the right external jugular vein following administration of 5 mg/kg Carprofen and 2 mg/kg 0.1% Bupivacaine analgesia and Avertin (2,2,2-tribromoethanol) anesthesia as described in detail by Thomsen and Caine (2007). Briefly, the catheter was inserted ∼12 mm into the jugular vein and anchored with sutures. The catheter was run subcutaneously to a mid-scapular incision and externalized using a single channel vascular access button (VAB62SMBS/25, Instech Laboratories, Plymouth Meeting, PA). Following surgery, mice were individually housed without enrichment for the duration of the experiment. Mice were allowed to recover for a minimum of 10 days before any testing began. Mice were tested daily in two-hour sessions at the same time each day during the light cycle.

### 2.2. Cocaine IVSA procedure

IVSA was performed using Med Associates (St. Albans, VT) operant conditioning chambers (307W) each enclosed in a sound attenuating cubicle (ENV-022MD). Two retractable response levers (ENV-310W) were mounted on the front wall of each chamber. A stimulus light (ENV-321W) was mounted above each lever. A house light (ENV-315W) with bulb (Chicago Miniature Lighting, LLC; CM1829) was centrally mounted on the rear wall of each chamber. An infusion pump (PHM-100;.05 RPM) was mounted inside the sound attenuating cubicle outside of the operant conditioning chamber. Chambers were controlled by Med Associates control units using MED-PC IV software (RRID:SCR_012156). Operant conditioning programs were written using MEDState notation and can be found at https://github.com/TheJacksonLaboratory/CSNA/tree/master/analysis/IVSA. Infusion supplies were provided by Instech Laboratories (Plymouth Meeting, PA) unless indicated otherwise. A 25-gauge single-channel plastic swivel (375/25PS) was mounted to a counterbalanced lever arm (SMCLA/MED) attached to the top of the chamber. Polyethylene tubing (BTPE-25) was used to connect a 60 ml plastic syringe (Becton Dickinson, Franklin Lakes, NJ) mounted in the infusion pump to the swivel, and to connect the swivel to the catheter port.

Cocaine hydrochloride was obtained from NIDA Drug Supply and cocaine doses were calculated as the salt. A single 10mg/kg dose of Methohexital (Brevital) was administered at the conclusion of IVSA testing to evaluate catheter patency. A daily 22.7 mg/kg antibiotic treatment of Enrofloxacin (Baytril) was administered throughout the experiment. Methohexital and enrofloxacin were obtained from Henry Schein, Inc. (Melville, NY). Methohexital and cocaine hydrochloride were dissolved in 0.9% saline. All solutions were filtered through 0.22 µm syringe filters.

Acquisition sessions used a fixed-ratio 1 (FR1) schedule of reinforcement at a dose of 1.0 mg/kg/infusion.

Each session began with illumination of the house light and extension of the two response levers. A left (active) lever press resulted in cocaine infusion and illumination of both stimulus lights for five seconds. This was followed by a twenty-second time-out during which the house light was off, and lever presses were recorded but had no consequences. Throughout the entire session, right (inactive) lever presses were recorded but had no consequences. The infusion pump delivered 9.03 µl/s of cocaine solution when engaged. Mice were weighed weekly, and the cocaine dose was adjusted to body weight by varying infusion time (100 ms/g). Acquisition criteria were met when 10 or more infusions were recorded for five sessions. Stabilization was defined as two consecutive sessions during which infusions did not vary by more than 20%. Thus, acquisition and stabilization require a minimum of five days. Mice that did failed to meet acquisition criteria within 28 training sessions were not tested further. For data analyses, we analyzed the number of sessions to meet acquisition criteria with a cut-off value of 28 sessions. If mice were advanced to the next stage of the IVSA protocol without reaching acquisition criteria, they were excluded from the ‘session to acquisition’ analyses. Additionally, for data analysis, we used the mean number of infusions received at FR1: 1.0 mg/kg/infusion after stabilization, and if mice did not reach acquisition or stabilization criteria, we recorded the number of infusions averaged over the entire acquisition period. Mice that became sick or injured, or lost catheter patency, were excluded from all acquisition analyses and did not advance to the dose response stage.

Dose-response sessions followed stabilization on the 1.0 mg/kg/infusion dose. For founders and J:DO mice, an eight-point dose response test was conducted with the following doses and in the following order: 1.0, 0.56, 0.32, 0.18, 0.1, 0.056, 0.032, 1.8 mg/kg/infusion. For all CC strains, a four-dose response test was conducted with the following doses and order: 1.0, 0.32, 0.1, 0.032, 1.0 mg/kg/infusion. The dose response curve was quantified by calculating an area under the curve (AUC) using the trapezoidal rule. Values for each mouse at each dose were calculated as the mean of the final two sessions of testing if the mice met stabilization criteria or the mean of all five sessions if mice did not meet the criteria. For data analyses, we compared AUC across strains and sexes, as well as infusions at each dose across the strains using a repeated measures approach.

Extinction sessions followed stabilization on the final session of the dose response stage. Mice were tethered as they were in all other stages; however, active lever presses had no consequence (i.e., house light remained on, stimulus lights were not illuminated, infusion pump was not activated) and no drug was delivered. Founder mice were tested for seven sessions under extinction conditions before advancing to the cued reinstatement stage; CC and J:DO mice were advanced to cued reinstatement when (1) the number of active lever presses decreased to 50% or less than the number of active lever presses on the first day of the extinction, and the difference between the final two sessions was less than 20% of the number of active lever presses on the first day, or (2) active lever presses were fewer than ten presses on or after day 3 of the extinction stage. CC and J:DO mice were therefore tested under these conditions for at least 3 days and a maximum of 9 days before advancing to reinstatement. For data analyses, we used the number of active lever presses across extinction sessions as repeated measures and also calculated the extinction rate as the negative slope of active lever presses across sessions.

Cue-induced reinstatement sessions followed the final extinction sessions. The sensory stimuli that were previously paired with cocaine infusions were delivered following an active lever press (i.e., infusion pump was turned on, house light was extinguished, cue lights were illuminated), but neither cocaine nor vehicle were infused. Mice were tethered, and the saline syringe was connected but the syringe was not placed in the infusion pump. For analyses of the reinstatement data, we excluded any mice that did not show a positive extinction rate (see Table 1). We then subtracted the number of active lever presses on the first day of reinstatement from the number of active lever presses on the final day of extinction; therefore, if a mouse increased lever pressing when moving from extinction to reinstatement, this value was positive.

Before each daily testing session, catheters were flushed with 20 µl of heparinized saline solution to ensure that catheters were clear of obstructions. Following each daily testing session, mice were infused with 2 µl/g enrofloxacin/saline solution (22.7 mg/kg) to prevent bacterial infection, and catheters were filled with 20 µl of heparin solution (100 U/ml heparin/saline) to maintain patency. At the end of the IVSA experiment, catheters were tested for patency with an100 µl infusion of a methohexital/saline solution (5 mg/kg). Rapid loss of muscle tone indicated patency. 97% of mice retained catheter patency at the conclusion of the testing period, including mice that did not reach acquisition criteria. Mice that had non-patent catheters were excluded from all data analyses.

### 2.3 Choice procedure

To test whether mice with different levels of cocaine self-administration also exhibited comparable response for cocaine relative to natural rewards (i.e., food), a choice test (Madsen & Ahmed, 2015) was employed. This multi-stage procedure consisted of shaping, food training, cocaine training, choice trials, and delayed choice trials. We compared two strains, CC002/UncJ that showed high cocaine IVSA intake, and CC003/UncJ that showed low intake. One operant wall had two active levers corresponding to the delivery of food reward (right) or a cocaine infusion (left). During training sessions, when a reinforcer was not available, its corresponding lever was not available. The single lever on the opposite wall was always inactive; responses are recorded without consequences.

The shaping phase establishes the basic contingency between lever presses and rewards and consisted of two days with one-hour shaping sessions. During these sessions, mice were placed into test chambers with the active lever and stimulus lights on. The right lever press (active lever) resulted in a palatable chocolate-flavored food reward (TestDiet, Catalog #1811142, 12.7% fat, 66.7% carbohydrate, and 20.6% protein). Pressing the active lever within a two-minute period led to illumination of the cue light, receipt of a food reward and the start of a 20-second timeout. During the timeout, the house light was extinguished, and any lever presses were recorded but had no consequence. If the period elapsed without a response, the cue lights were illuminated, and a food reward pellet was released into the magazine, followed by a timeout, ensuring that all mice experienced the reward and the associated cues, even if they did not initially press the lever.

Phase I of food training consisted of eight daily, one-hour sessions, during which mice needed to press the active lever to receive a food reward (no automatic delivery). Phase II of food training commenced after jugular vein catheterization and recovery. During this three-day phase, mice were placed into the operant chamber and the IVSA tether was connected to a saline-filled syringe in the infusion pump. Although the subjects did not receive any infusions during this phase, mice acclimated to the tether connection. Mice had to press the active lever to receive food, reinforcing the association of active lever press and food reward delivery.

Mice were trained to self-administer cocaine during three daily, two-hour sessions, on a fixed-ratio 1 (FR1) schedule 0.32 mg/kg/infusion following the IVSA acquisition procedure described above. This dose elicited the highest response rate in the founder dose-response study. The infusion pump delivered 9.03 µl/s of cocaine solution when engaged. Mice were weighed weekly, and the cocaine dose was adjusted for body weight by varying infusion time (100 ms/g).

Following three days of cocaine IVSA training, a one-day choice test assessed preference between drug and natural reward (i.e., food). Choice sessions used the FR1 schedule and 0.32 mg/kg/infusion of cocaine or 1 food pellet. During this two-hour session, consisting of 15 eight-minute trials, both active levers and associated stimulus lights were available to allow mice to select food, drug, or neither. If a mouse chose either the food or cocaine lever, a timeout occurred in which stimulus lights were extinguished and the mouse received its reward. If a mouse did not press either active lever for six minutes during a trial, the trial was recorded as an omission; in response, a timeout occurred, and the mouse moved to the next eight-minute trial. After the one-day choice trial, mice underwent four repeating four-day choice cycles, a total of 16 sessions. During these cycles, IVSA sessions occurred on Days 1, 2, and 3, and a choice trial occurred on Day 4. This resulted in a total of 12 cocaine IVSA sessions and 4 choice trials. For the next eight days, mice underwent daily two-hour delayed-choice trials in which there was an increasing delay between each food lever press and the pellet delivery. Every two days, the length of delay increased (15 sec, 30 sec, 45 sec, 60 sec).

### 2.4 Punishment procedure

To assess how the chance of an adverse consequence (foot shock) influenced drug-seeking behavior, we evaluated different subjects from the same two strains that were tested in the Choice procedure, CC002/UncJ (high cocaine intake) and CC003/UncJ (low cocaine intake), plus an additional founder strain with reliable cocaine acquisition during IVSA, NZO/HILtJ. After completing the cocaine IVSA acquisition procedure described above, mice began a seven-day testing regimen. All sessions used an FR1 schedule at a dose of 0.32 mg/kg/infusion at 9.03 µl/s. Cocaine punishment sessions lasted two hours each day.

Each session began with illumination of the house light and extension of the two response levers. Pressing the active lever resulted in a cocaine infusion and illumination of the stimulus light for five seconds. Each press on the active lever was associated with a 50% chance of receiving a contingent 500 ms foot shock. This was followed by a twenty-second timeout during which the house light was extinguished, and lever presses were recorded but had no consequences. Throughout the entire session, presses on the inactive lever were recorded but had no programmed consequences, ensuring that any lever pressing behavior was specifically directed towards obtaining cocaine despite the 50% probability of foot shock. The cocaine dose was adjusted weekly according to body weight by varying the infusion pump time (100 ms/g).

Each testing day, the shock intensity increased by 0.05 mA, starting at 0mA and reaching 0.35 mA on day seven. This enabled observation of how escalating punishment influenced lever pressing for cocaine. The number of active and inactive lever presses were recorded in each session, providing data on the behavioral response to the probability of punishment. Additionally, the number of infusions and the time (in seconds) between a foot shock and the next active lever press were collected to construct a shock-response curve, allowing for a detailed analysis of how punishment affected drug-seeking behavior.

### 2.5 Statistical methods

All results were computed in the R statistical computing environment (Version 4.4.3; R Core Team, 2024). We assessed effects of strain and sex independently and strain by sex interaction using separate two-way ANOVAs for each IVSA trait for the founder and CC strains using the base R package, stats (RRID:SCR_0259680), and the ‘lm’ and ‘anova’ functions. For dose response and extinction, we also conducted two-way repeated measures ANOVAs to test for strain differences over the course of each stage for the founder and CC strains using the R package, afex (RRID:SCR_0228570), and the function ‘aov_car’ with ‘subject’ as random effect, and ‘sessions’ and ‘strain’ as factors. To address potential non-normality, we performed a residual analysis on the models by plotting the fitted values against the residuals to assess model fit. We applied appropriate transformations to improve normality of each dataset then fitted a second ANOVA using the transformed variable. Specifically, for the two-way ANOVAs, we did not transform the sessions to acquisition data for either the founders or CC strains, we used a square root transformation for infusions at stabilization for both founders and CC strains, performed no transformation on the AUC founder data but used a logp1 transformation for the AUC data in the CC strains, and performed z-rank transformations for extinction rate and the change in active lever presses from the last extinction session to the first reinstatement session for both the founder and CC strains.

For each IVSA trait analyzed in the founder and the CC strains, we calculated heritability using a linear modeling approach as previously described (Saul et al., 2020). Briefly, the individual inbred strains were treated as a categorical variable in the following equation:

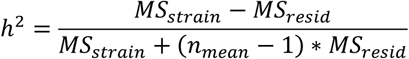

where *MS_strain_* is the mean square of the strain effect, *n_mean_* is the mean number of samples within each strain, and *MS_resid_* is the mean square of the residuals. For each trait, we used the same transformations as implemented in the two-way ANOVAs (see above). Linear mixed models were built using the ‘lmer’ function from the lme4 package (RRID:SCR_015654) in R. Models included genotype (strain) as a random effect. After fitting the model using REML, between and within-group variance were extracted to calculate heritability.

For each IVSA trait analyzed above in the CC strains we calculated the strain mean and used the ‘corr.test’ function in the ‘psych’ package in R to perform Spearman’s rank correlation of all complete pairwise comparisons. For example, if a strain did not meet acquisition criteria, there would be no IVSA trait associated with extinction and reinstatement stages, therefore these strains were excluded from the pairwise correlations calculations just for those IVSA stages. To control for multiple testing using the false discovery rate (Benjamini & Hochberg, 1995), we calculated adjusted p-values using ‘p.adjust’ in the ‘stats’ R package, and selected the ‘fdr’ method.

In the IVSA choice experiment, mice could press levers for food or cocaine or not press any lever, which was counted as an omission. When there was no delay for the choice trial, we tested whether the strains exhibited different mean proportion of lever presses for cocaine over all trials with Welch’s one-sided *t*-test for unequal variances. When the choice test included a delay, we fit the response with a Poisson mixed-effect model where the response was the total number of lever presses for cocaine over the two sessions per delay (in seconds), with subject as the random effect, strain as a fixed effect and an offset term as the logarithm of the total trials. Accounting for strain as a fixed effect, we tested whether there was a strain by delay interaction with the likelihood ratio test where the restricted model included strain and the random effect for subject and the full model included strain, the random effect for subject, and the interaction term for strain and delay.

In the IVSA punishment experiment, we recorded and analyzed the duration (in seconds) between an active lever press with a foot shock and the next active lever press. Each subject is exposed to the active lever at each shock intensity. To account for the correlation of repeated measures, we fit a mixed-effect linear model with random effects for subject and subject crossed with shock intensity, including varying intercepts and varying slopes (Doran, 2007). Natural log transformation was used stabilize the variance in time to the next active lever press after a foot shock. After accounting for random effects of subject, subject crossed with shock intensity, and fixed effects of strain and shock intensity, we tested whether there was a strain by shock intensity interaction with the likelihood ratio test where the restricted model included the random effects and fixed effects strain and shock intensity, and the full model included all parameters in the restricted model and the strain by shock intensity interaction. Restricted and full models were fit using the R package ‘lme4’ (Bates, 2015) with REML set to false. Mixed effect model estimates and Wald 95% confidence intervals for Figure 7 were calculated on the full model with the R package ‘broom.mixed’ (RRID:SCR_026712; Bolker, 2024) and visualized using the R package ‘ggplot2’ (Wickham, 2016).

## 3. Results

### 3.1 Cocaine IVSA

Most founder, CC and J:DO mice reached acquisition criteria. However, no 129S1/SvImJ founder strain mice or mice from three CC strains (CC010/GeniUncJ, CC037/TauUncJ, CC044/UncJ) reached acquisition criteria (Table 1). Analysis of the number of sessions to reach acquisition criteria in the founders revealed an effect of strain (F_7, 178_ = 23.72; p < 0.001; Table 2) and an effect of sex (F_1, 178_ = 4.28; p = 0.040) with female acquiring slower than males (Figure S1), but no strain by sex interaction effect (F_7, 178_ = 0.41; p = 0.893). The analysis of sessions to acquisition in the CC strains revealed an effect of strain (F_49, 278_ = 8.34; p < 0.001) but no sex effect (F_1, 278_ = 0.25; p = 0.694) or strain by sex interaction (F_48, 278_ = 1.15; p = 0.241. The average number of infusions per day, considering only the two days when the mice met stabilization criteria, or over the entire acquisition stage for those that did not meet stabilization criteria, is a measure of cocaine intake. Analysis of mean infusions in founders by two-way ANOVA with strain and sex as the independent variables revealed an effect of strain (F_7, 179_ = 35.05; p < 0.001) but not an effect of sex (F_1, 179_ = 0.16; p = 0.724) or a strain by sex interaction (F_7, 179_ = 0.47; p = 0.850). Similarly in the CC strains, analysis of mean infusions revealed an effect of strain (F_49, 283_ = 5.78; p < 0.001) but not an effect of sex (F_1, 283_ = 0.29; p = 0.588) or a strain by sex interaction (F_48, 283_ = 1.01; p = 0.455). The mean infusions at stabilization ranged widely by strain (Figure 1B). Among the founder strains, PWK/PhJ recorded the highest (followed by CAST/EiJ) and 129S1/SvImJ the lowest (exceeded slightly by WSB/EiJ). Among the CC strains, the mean infusions ranged to and beyond those recorded for founder strains, with two CC strains recording more infusions than PWK/PhJ and five CC strains falling below WSB/EiJ (Figure 1B). Individual J:DO mice displayed a wide range of responses, encompassing the full range of values observed across all founder strains (Figure 1C).

**Figure 1.**
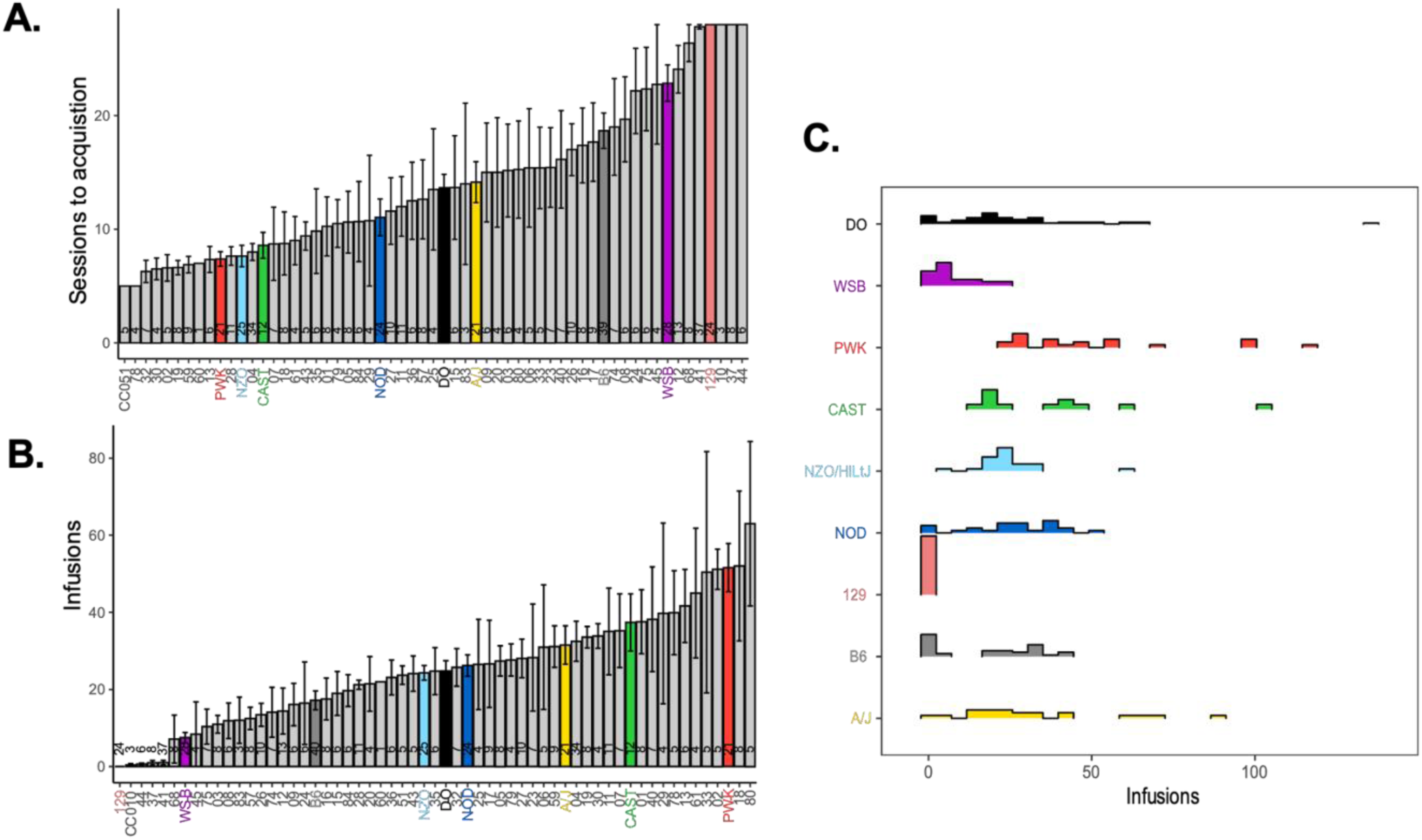
Cocaine IVSA acquisition and infusions at stabilization by strain. (A) Mean ± standard error (SE) of the number of sessions until acquisition criteria were met for each strain. Note, if acquisition criteria were never met, the maximum number of sessions that mice were exposed to the IVSA paradigm (28 sessions) are reported. (B) Mean ± SE infusions (1mg/kg dose) during the two-day stabilization period after acquisition criteria is met. (C) Frequency distribution of number of infusions (1mg/kg dose) in J:DO and founder strains. In all panels, CC strains are denoted in grey, J:DO population in black, and founder strains in color.

**Table 2.** Results from ANOVA tests in founder and CC strains across IVSA stages.

| Stage | Trait | ANOVA | Predictor | Founder strains |  |  | CC strains |  |  |
| --- | --- | --- | --- | --- | --- | --- | --- | --- | --- |
|  |  |  |  | F | df | p | F | df | p |
| Acquisition | Sessions to acquisition | 2-way | Strain | 23.72 | 7,178 | **<0.001** | 8.34 | 49,278 | **<0.001** |
|  |  |  | Sex | 4.28 | 1,178 | **0.040** | 0.25 | 1,278 | 0.694 |
|  |  |  | Interaction | 0.41 | 7,178 | 0.893 | 1.15 | 48,278 | 0.241 |
| Acquisition | Infusions at stabilization | 2-way | Strain | 35.05 | 7,179 | **<0.001** | 5.78 | 49,283 | **<0.001** |
|  |  |  | Sex | 0.16 | 1,179 | 0.724 | 0.29 | 1,283 | 0.588 |
|  |  |  | Interaction | 0.47 | 7,179 | 0.850 | 1.01 | 48,283 | 0.455 |
| Dose Response | Infusions | 2-way repeated measures | Strain | 8.65 | 6,94 | **<0.001** | 8.47 | 48,258 | **<0.001** |
|  |  |  | Dose | 63.30 | 7,658 | **<0.001** | 118.85 | 3,774 | **<0.001** |
|  |  |  | Interaction | 1.27 | 42,658 | 0.118 | 2.87 | 144,774 | **<0.001** |
| Dose Response | AUC | 2-way | Strain | 8.92 | 6,78 | **<0.001** | 4.56 | 45,142 | **<0.001** |
|  |  |  | Sex | 1.38 | 1,78 | 0.243 | 4.53 | 1,142 | **0.035** |
|  |  |  | Interaction | 3.39 | 6,78 | **0.005** | 1.00 | 42,142 | 0.477 |
| Extinction | Rate | 2-way | Strain | 2.70 | 6,78 | **0.019** | 42.59 | 45,139 | 0.290 |
|  |  |  | Sex | 0.32 | 1,78 | 0.539 | 3.42 | 1,139 | **0.045** |
|  |  |  | Interaction | 4.20 | 6,78 | 0.554 | 56.39 | 42,139 | **0.022** |
| Extinction | Active lever presses | 2-way repeated measures | Strain | 6.20 | 6,79 | **<0.001** | 5.06 | 48,200 | **<0.001** |
|  |  |  | Session | 12.62 | 6,474 | **<0.001** | 14.23 | 2,400 | **<0.001** |
|  |  |  | Interaction | 1.20 | 36,474 | 0.202 | 0.89 | 96,400 | 0.754 |
| Reinstatement | Change in active lever presses | 2-way | Strain | 2.90 | 6,67 | 0.815 | 59.18 | 45,81 | **0.023** |
|  |  |  | Sex | 0.41 | 1,67 | 0.521 | 1.40 | 1,81 | 0.187 |
|  |  |  | Interaction | 3.92 | 6,67 | 0.660 | 32.51 | 38,81 | 0.378 |

Bolded text indicates p-values < 0.05.

To assess the relative sensitivity to the reinforcing effects of cocaine infusions at different doses, we analyzed the number of infusions during dose response using a repeated measures ANOVA for founder strains, which revealed an effect of strain (F_6, 94_ = 8.65; p < 0.001) and dose (F_7, 658_ = 63.30; p < 0.001) but no strain by dose interaction (F_42, 658_ = 1.27; p = 0.118). Analysis of CC strain data also showed an effect of strain (F_48, 258_ = 8.47; p < 0.001) and dose (F_3, 774_ = 118.85; p < 0.001), as well as a strain by dose interaction (F_144, 774_ = 2.87; p < 0.001). We then calculated the area under the curve (AUC) as a composite measure of dose responsiveness for CC strains (Figure 2B) and founders (Figure 2C, 2D). Analysis of the AUC for founders revealed an effect of strain (F_6, 78_ = 8.92; p < 0.001) and a strain by sex interaction (F_6, 78_ = 3.39; p = 0.005; Figure S2), but not a sex effect (F_1, 78_ = 1.38; p = 0.243). Analysis of the AUC for CC strains revealed a main effect of strain (F_45, 142_ = 4.56; p < 0.001) and sex (F_1, 142_ = 4.53; p = 0.035) with females exhibiting a greater AUC than males (Figure S3), but not a strain by sex interaction (F_42, 142_ = 1.00; p = 0.477). The dose-response AUC for the J:DO population was also highly variable (Figure 2A), and the founder strains PWK/PhJ and WSB/EiJ displayed distinct drug use preferences, as PWK/PhJ used large amounts of cocaine and WSB/EiJ used very little cocaine across the dose response curve (Figure 2C, 2D).

**Figure 2.**
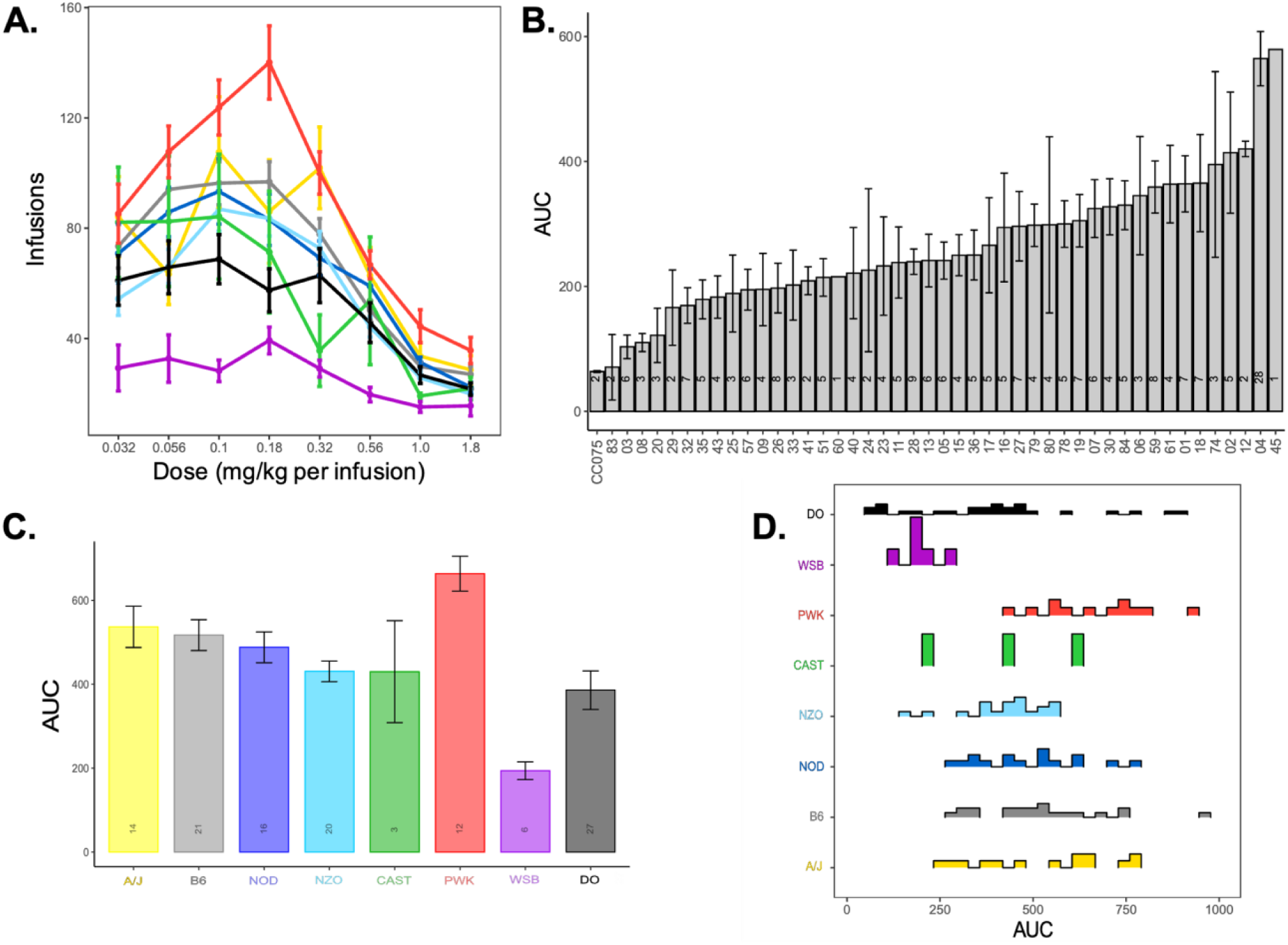
Cocaine IVSA dose response by strain. (A) Mean ± SE infusions at various doses offered during a two hour IVSA session in the founder strains; color identifies strain as labeled in other panels of figure. (B) Mean ± SE area under the curve (AUC) calculated across multiple dose categories in founders (B) and CC strains (C). (D) Frequency distribution of AUC in J:DO and founder strains. In all panels, CC strains are denoted in light grey, J:DO population in dark grey or black, and founder strains in color.

Analysis of the number of active lever presses over the seven-day extinction period for founder strains by repeated measures ANOVA showed an effect of strain (F_6, 79_ = 6.20; p < 0.001; Figure 3) and session (F_6, 474_ = 12.62; p < 0.001), but not a strain by session interaction (F_36, 474_ = 1.20; p = 0.202). Analysis of CC strains conducted using the first 3 days of extinction due to the criteria based 3-to-9-day extinction protocol revealed an effect of strain (F_48,200_ = 5.06; p < 0.001) and session (F_2, 400_ = 14.23; p < 0.001) but not a strain by session interaction (F_96, 400_ = 0.89; p = 0.754). The analysis of the rate of extinction in the founders revealed a main effect of strain (F_6, 78_ = 2.70; p = 0.019), but not an effect of sex (F_1, 78_ = 0.32; p = 0.539) or strain by sex interaction (F_6, 78_ = 4.20; p = 0.554). While the similar analysis in the CC strains revealed an effect of sex with males (F_1, 139_ = 3.42; p = 0.045) showing a greater rate of extinction than females (Figure S4), and a strain by sex interaction (F_42, 139_ = 56.39; p = 0.022), yet not a main effect of strain (F_45, 139_ = 42.59; p = 0.290; Figure 4). The distribution of the extinction rates in J:DO mice encompasses the full range of values observed across all founder strains, among which, NOD/ShiLtJ strain displayed the greatest rate of extinction and WSB/EiJ the lowest rate.

**Figure 3.**
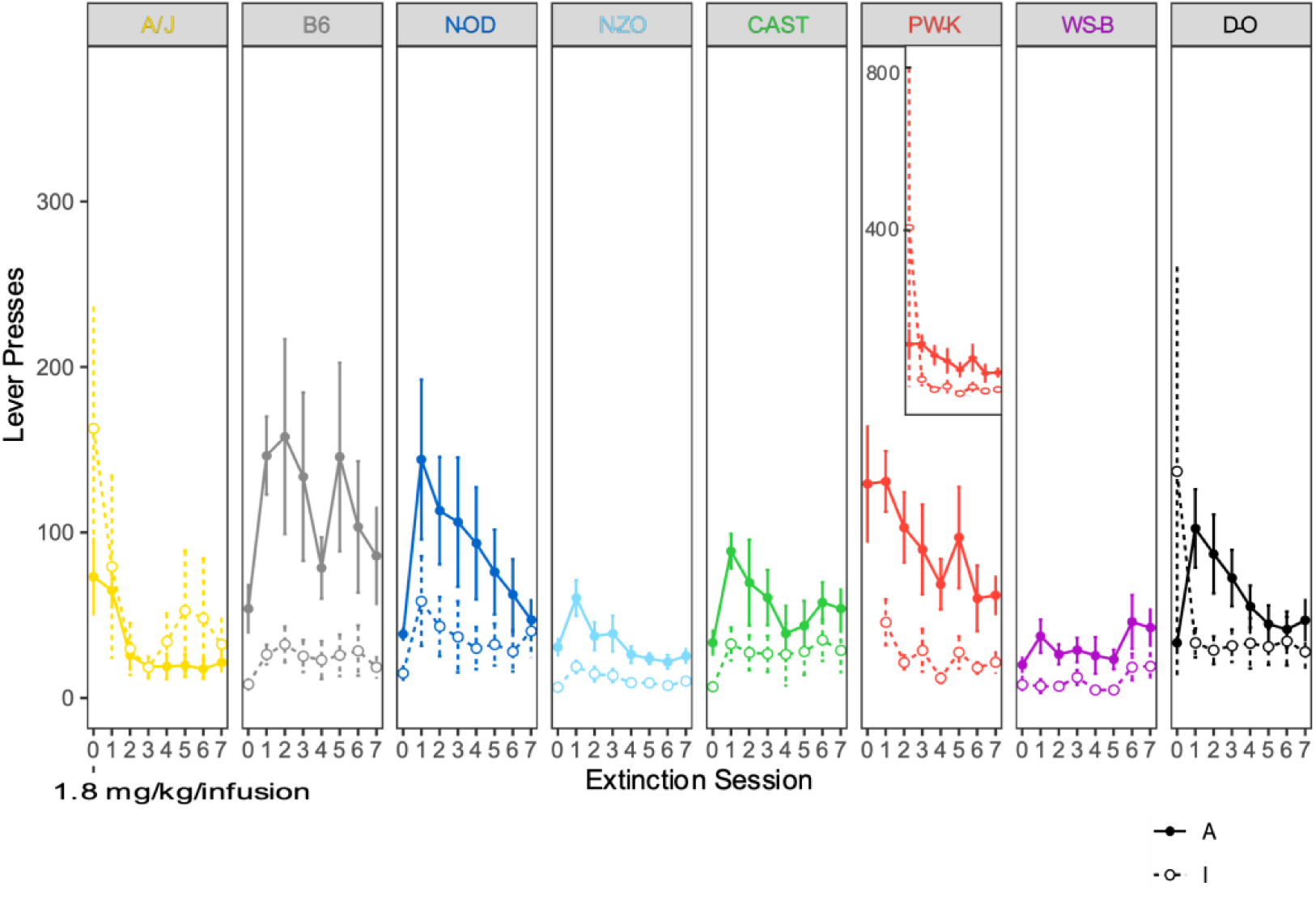
Extinction of cocaine IVSA behavior in founder strains. Mean ± SE of active (A; filled circle and solid line) and inactive (I; open circle and dashed line) lever presses for each founder strain following the final day of dose response testing at 1.8 mg/kg dose and for seven subsequent testing sessions where no cocaine was available.

**Figure 4.**
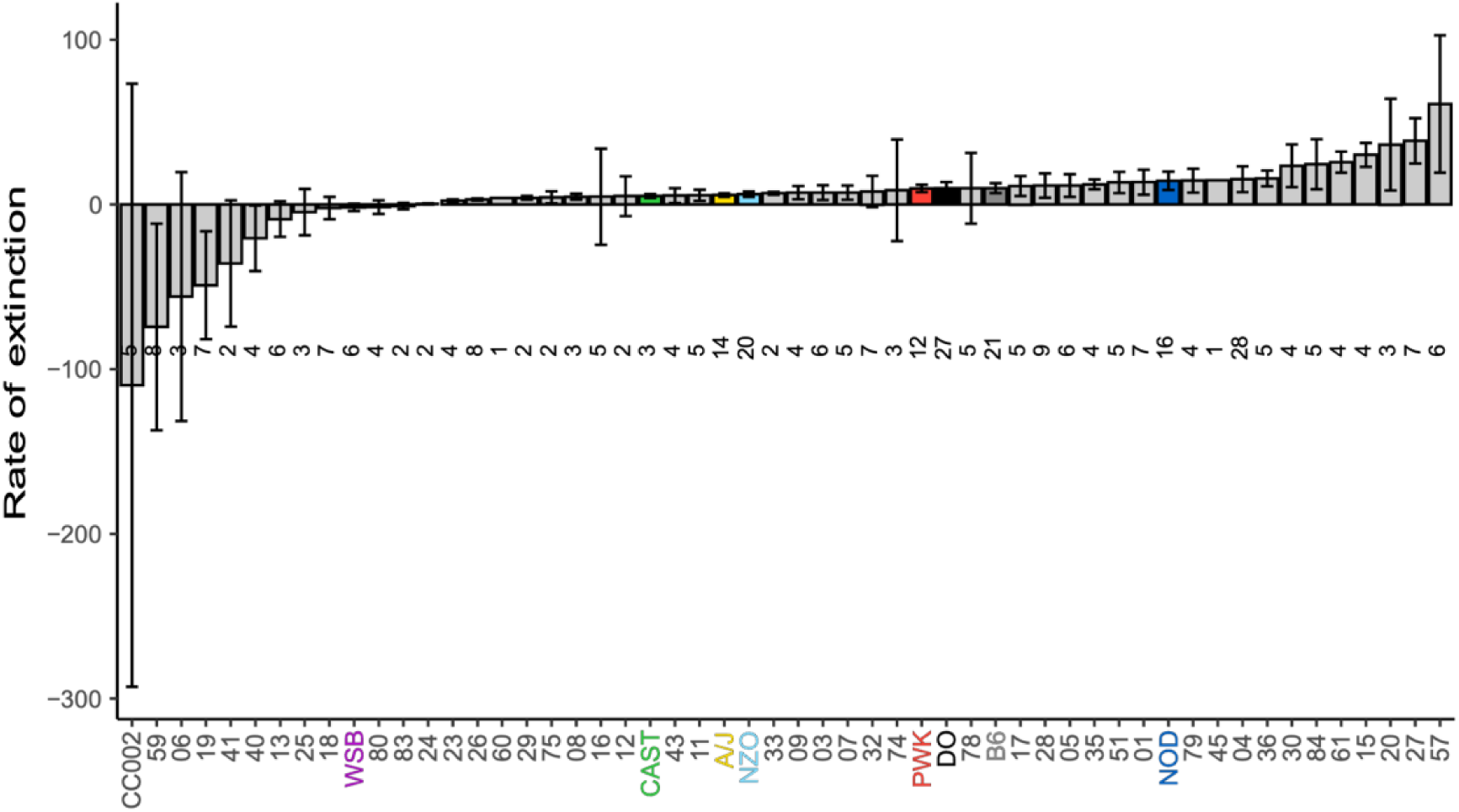
Cocaine IVSA extinction by strain. Mean ± SE rate of extinction representing change in active lever pressing across extinction sessions. CC strains are denoted in grey, J:DO population in black, and founder strains in color.

After restricting the data analysis to include only mice with a negative slope of extinction, which included mice from all strains that had reached acquisition criteria (Table 1), we analyzed the change in active lever presses between the final day of extinction and the first day of cued reinstatement. In the founder strains, the analysis showed no effect of strain (F_6, 67_ = 2.90; p = 0.815; Figure 5), sex (F_1, 67_ = 0.41; p = 0.521), or a strain by sex interaction (F_6, 67_ = 3.92; p = 0.660). However, analysis of this trait in CC strains did reveal an effect of strain (F _45, 81_ = 59.18; p = 0.023; Figure 5), yet not an effect of sex (F_1, 81_ = 1.40; p = 0.187 or a strain by sex interaction (F_38, 81_ = 32.51; p = 0.378). The distribution of the J:DO response encompasses the full range of values observed across all founder strains, among which, NOD/ShiLtJ and CAST/EiJ strains displayed the greatest cued-reinstatement response and WSB/EiJ the lowest (Figure 5).

**Figure 5.**
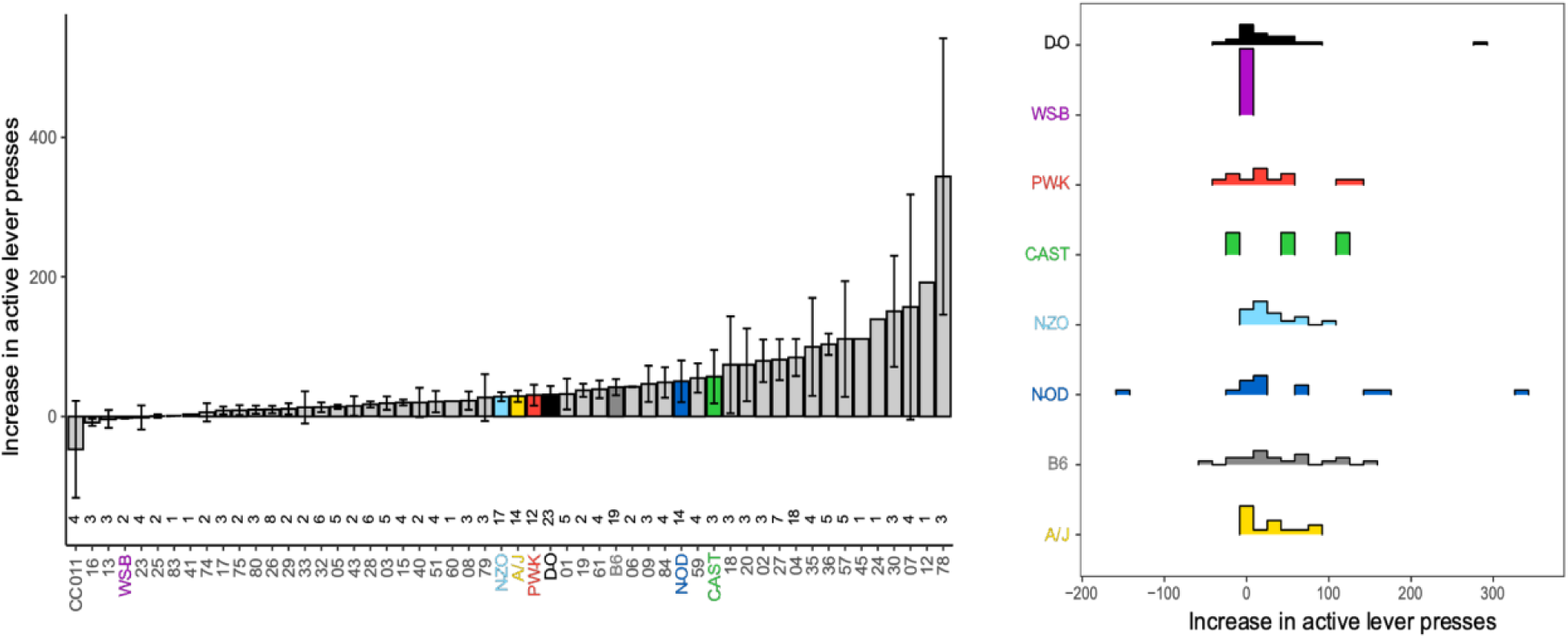
Reinstatement of cocaine IVSA behavior following extinction by strain. (A) Mean ± SE percent (%) increase in active lever presses during reinstatement compared to presses on the final extinction session. (B) Frequency distribution of % increase in active lever presses during reinstatement compared to presses on the final extinction session. In all panels, CC strains are denoted in grey, J:DO population in black, and founder strains in color.

The heritability of each IVSA trait ranged in founders from 0 to 0.585, while in the CC strains heritability ranged from 0.019 to 0.421 (Table 3). While we found that some IVSA traits from different stages are correlated with one another in the CC strains, not all are (Table 3). The number of sessions to meet acquisition criteria is negatively correlated with the mean number of infusions at stabilization (ρ =-0.773, df = 57, p < 0.001), and the change in active lever presses from the last day of extinction to the first day of reinstatement is positively correlated with dose response AUC (ρ = 0.436, df = 52, p = 0.009). Moreover, the AUC is positively correlated with infusions at stabilization (ρ = 0.380, df = 52, p = 0.032), yet after adjusting for multiple testing, we find a slightly weaker relationship (p = 0.061). Similarly, the change in active lever presses from the last day of extinction to the first day of reinstatement is positively correlated with the rate of extinction (ρ = 0.388, df = 52, p = 0.030), but after adjusting for multiple testing, there is a weaker relationship (p = 0.061).

**Table 3.** IVSA trait heritability estimates (h^2^) for founder and CC strains, and correlations results within CC strains.

| IVSA trait | Founder $h^2$ | CC $h^2$ | CC correlation: Sessions to acquisition | | | CC correlation: Infusions at stabilization | | | CC correlation: Dose response AUC | | | CC correlation: Rate of extinction | | |
| --- | --- | --- | --- | --- | --- | --- | --- | --- | --- | --- | --- | --- | --- | --- |
| | | | $\rho$ | p | adj-p | $\rho$ | p | adj-p | $\rho$ | p | adj-p | $\rho$ | p | adj-p |
| Sessions to acquisition | 0.534 | 0.421 | - | - | - | - | - | - | - | - | - | - | - | - |
| Infusions at stabilization | 0.585 | 0.327 | -0.670 | **<0.001** | **<0.001** | - | - | - | - | - | - | - | - | - |
| Dose response AUC | 0.435 | 0.365 | -0.201 | 0.727 | 0.865 | 0.380 | **0.032** | 0.061 | - | - | - | - | - | - |
| Rate of extinction | 0.141 | 0.019 | -0.150 | 0.933 | 0.933 | -0.107 | 0.933 | 0.933 | 0.165 | 0.933 | 0.933 | - | - | - |
| Change in active lever presses from extinction to reinstatement | 0.000 | 0.146 | -0.275 | 0.263 | 0.387 | 0.124 | 0.933 | 0.933 | 0.436 | **0.009** | **0.020** | 0.388 | **0.030** | 0.061 |

Bolded text indicates p-values or adjusted p-values < 0.05.

### 3.2 Cocaine IVSA or food choice test and delayed choice test

When there was no delay in reward, CC002/UncJ mice, shown to have high self-administration in the acquisition study, chose cocaine (cocaine choice average = 29.6%) more frequently than a strain shown to have lower levels of self-administration, CC003/UncJ mice (cocaine choice mean = 6.2%), with a 95% lower bound of 12.7% more than CC003/UncJ mice (t = 3.77, df = 22.16, p < 0.001). When a delay in reward was introduced to lower the salience of food reward, accounting for the fixed effect of strain, we found an interaction between strain and delay (*X*_1_^2^ = 9.5, p = 0.002), such that CC003/UncJ mice reduced their response, whereas in CC002/UncJ, the mice maintained a high level of response (Figure 6).

**Figure 6.**
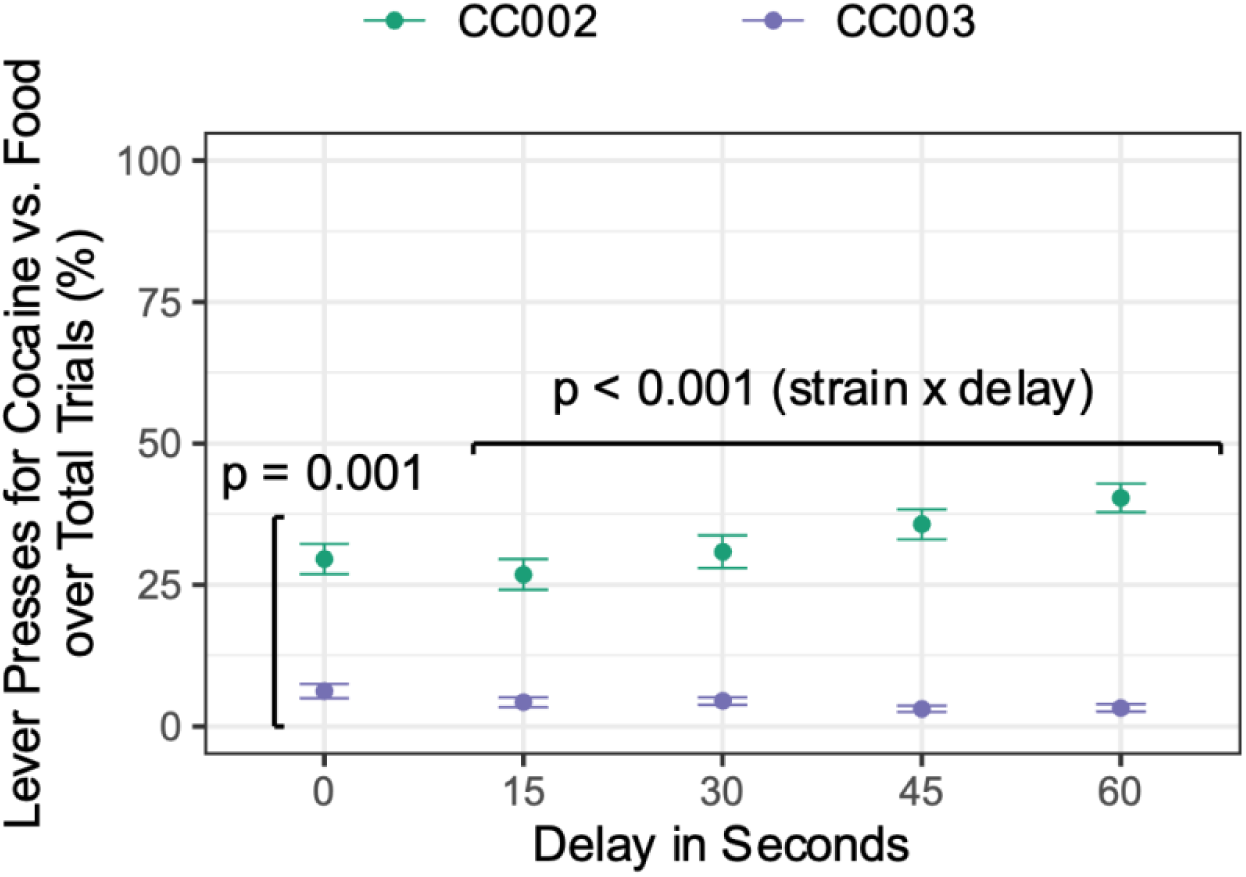
IVSA cocaine vs food choice. Mean ± SE number of active lever presses for cocaine versus food over total trials (%) with no delay (delay in seconds = 0) and after a delay (14, 30, 45, and 60 seconds) is introduced prior to the reward in CC002/UncJ (green) and CC003/UncJ (purple) strains.

**Figure 7.**
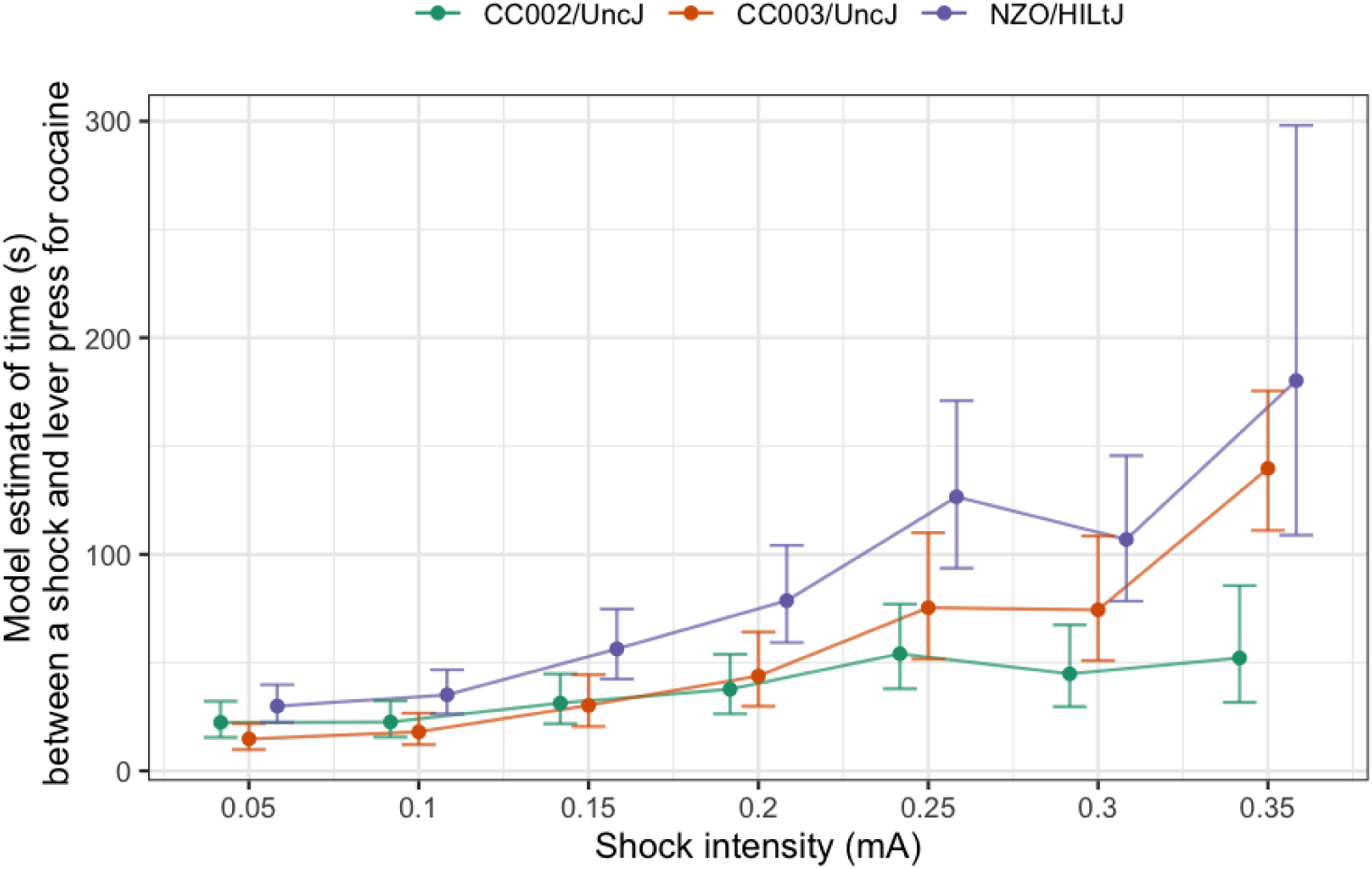
IVSA cocaine with punishment. Mixed-effect model estimated strain means with 95% Wald confidence intervals at each shock intensity (mA) of time (in seconds) to an active lever press after a foot shock. Model estimates were calculated for each subject using the mixed effect model that included random effects for subject and subject crossed with shock intensity, fixed effects for strain and shock intensity and the strain by shock intensity interaction effect. CC002/UncJ (green), CC003/UncJ (orange), and NZO/HILtJ (purple) estimated strain means and 95% confidence intervals are shown in the figure with points jiggered on the x-axis to improve visualization.

### 3.3 Cocaine IVSA with punishment

We compared CC002/UncJ and CC003/UncJ in the effect of shock intensity on the delay between punishment and the next active lever press using the likelihood ratio test on mixed-effect models. In a model including fixed effects of strain and shock intensity, and random effects of subject and subject crossed with strain intensity, we found a strain by shock intensity interaction effect (*X*_1_^2^ = 32.9, p < 0.001) in the logarithm of the time between a foot shock and the next active lever press. As shock intensity increased, all strains took more time to press the active lever after foot shock, however CC002/UncJ was slower to change its behavior than CC003/UncJ and NZO/HILtJ strains, both of which more rapidly increased their delay in responding (Figure 7).

## 4. Discussion

Here we demonstrate significant strain and sex differences across multiple key stages of cocaine IVSA response, including acquisition, dose-response, extinction, and cue-induced reinstatement of self-administration behavior, with each stage modeling different aspects of vulnerability to CUD. Among the CC/J:DO founders, the five classical laboratory strains (C57BL/6J, A/J, 129S1/SvImJ, NOD/ShiLtJ, and NZO/HILtJ) and three wild-derived strains (CAST/EiJ, PWK/PhJ, and WSB/EiJ) display highly disparate phenotypic responses across all stages, indicating the presence of variation in genetic influences on initiation of cocaine self-administration and the subsequent maintenance of these responses. Most notably, the widely used C57BL/6J strain, and the wealth of mutant and gene-deletion strains derived from it, is frequently used to study the biological mechanisms and behaviors related to cocaine response, yet we found that C57BL/6J is not the most extreme on any measure we examined. We note that C57BL/6J exhibits volitional response for cocaine to some degree; however, the wild-derived strains CAST/EiJ and PWK/PhJ exhibit a higher rate of cocaine self-administration and reach acquisition criteria faster and thus may be better models of susceptibility to some features of CUD. This is consistent with reports of genetically fixed regions in the common inbred strains widely thought to be related to historical selection for docility and the subsequent restoration of behavioral variation in the CC strains (Chesler, 2014; Philip et al., 2011; Yang et al., 2007). The 44 CC strains and J:DO mice, which were derived from these eight founders, demonstrated an even wider range of phenotypic variation relative to the founder strains. This observed transgressive segregation suggests that these highly recombinant strains each harbor multiple cocaine self-administration vulnerability alleles. We also demonstrate that strains which vary in acquisition and drug intake during acquisition also show differences in more complex assays of addiction related behaviors.

We observed substantial heritability of multiple IVSA traits in the founder and CC strains indicating a strong genetic contribution and suitability for downstream genetic analyses. Moreover, J:DO mice displayed a high degree of inter-individual behavioral variation that, like CC stains, extend beyond the range of the founders, revealing the importance of heterozygosity and epistasis across loci. Finally, we show that some aspects of cocaine self-administration are correlated, particularly for traits within the same IVSA phase, for example strains that reach acquisition criteria in fewer sessions self-administer a greater number of infusions. Reinstatement and dose response behaviors are also significantly correlated indicating shared mechanisms of drug seeking as reward availability or salience changes. However, others such as extinction and reinstatement of cocaine self-administration are only somewhat correlated with one another. The differences observed among stages indicate that they are likely under distinct genetic regulation, highlighting the importance of multi-stage, longitudinal studies to identify stage-specific genes and variants underlying cocaine use.

We found strong, consistent, and significant strain differences in the number of sessions to reach acquisition and cocaine intake in the 8 founder and the 44 CC strains, consistent with previous reports of strain differences in cocaine self-administration in populations of rodents with more limited genetic diversity. Strain differences in IVSA acquisition have also been shown in rats (Kosten et al., 1997) and mice (Deroche et al., 1997; Kuzmin & Johansson, 2000; Roberts et al., 2018; Schoenrock et al., 2020; Thomsen & Caine, 2006), including among the BXD recombinant inbred population (Dickson et al., 2016), the hybrid mouse diversity panel (Bagley et al., 2022; Khan et al., 2023), and in a crossbreeding study using cocaine-preferring and non-preferring mouse strains (Ruiz-Durantez et al., 2006). Additionally, we show high heritability, with estimates for acquisition traits in the CC strains and their founders ranging between 0.327 and 0.585, which is greater than heritability for acquisition observed in the recombinant inbred BXD population (h^2^ = 0.28; Dickson et al., 2016).

Our results revealed an important sex difference in cocaine acquisition in the founder strains; males reach acquisition criteria in fewer sessions. Sex differences in cocaine self-administration have been observed in rats (Becker, 2016; Hu et al., 2004; Roth & Carroll, 2004), specifically, that females acquire cocaine IVSA faster than males (de Guglielmo et al., 2024; Lynch & Carroll, 1999), which contrasts with our results in the founder mouse strains. Earlier work in mice has shown that a heightened propensity to acquire cocaine self-administration in females compared to males is contingent on dose, reinforcement schedule, and estrous cycle phase (Griffin et al., 2007; Martini et al., 2015). Further, we found that in the CC strains, females exhibited a higher AUC of dose response curve than males, and in the founder strains, we found a significant strain by sex interaction for AUC. Strain-specific sex differences in the same behavior can be dependent or independent of estrous cycle effects across genetically diverse populations (Mogil et al., 2000), such that the effect may be developmental in some strains whereas in others it may be an effect of gonadal steroid cycling (McEwen & Milner, 2017).

Dose-response relationships, reflected in the AUC of the dose response curve reflect the relative sensitivity to the reinforcing effects of cocaine and predict sensitive and resistant phenotypes (Sizemore et al., 1997). Our results revealed significant strain effects on AUC in both founder and CC strains. The observed strain effects are consistent with previously reported differences in the cocaine dose-response curve for in the recombinant inbred BXD population (Dickson et al., 2016) and a smaller panel of eight mouse strains (Roberts et al., 2018), yet the latter study focused on males and our findings of sex difference in dose response highlight the importance of examining both sexes. The classic inverted-U shape of our dose response curve results from an increase in relative reinforcing effects of the drug as a function of increases in dose up to a certain point, beyond which a decline in rate of responding is observed, likely due to the adverse effects at high doses (Meisch & Stewart, 1995; Pickens & Thompson, 1968; Skjoldager et al., 1991). Subjects who are vulnerable are likely to initiate and maintain self-administration at low cocaine doses (left shifted dose-response curve) or continue to respond at a higher rate and consume higher quantities of the drug across doses (upward shifted dose-response curve; (Piazza et al., 2000). Our results reveal strains with dose response characteristics consistent with a susceptible phenotype, as exemplified by the upward shifted dose response curve for PWK/PhJ relative to WSB/EiJ. Finally, the heritability measures for the dose-response AUC were 0.435 in the founders and 0.365 in the CC mice, indicating considerable genetic influences over the responses generated by varying drug doses.

Responses during consecutive extinction sessions provide a measure of perseverative drug-seeking in the absence of response–contingent drug delivery (Markou et al., 1993). The inability to limit responses during extinction is interpreted as compulsive drug-seeking, yet we note that a brief extinction period does not prima facie recapitulate abstinence in long term SUD. We found significant founder strain effects in active lever presses during extinction, and in CC strains, we found this behavior also varies by sex with males extinguishing faster than females. In rats, extinction of cocaine self-administration is similar (Miguens et al., 2013) or diverges in a temporally specific manner (Kruzich & Xi, 2006) between Lewis and Fischer 344 inbred strains. Re-exposure to drug-associated cues after extinction can trigger reinstatement in laboratory animals as evidenced by increased active lever presses in the presence of drug paired cues (Fuchs et al., 2003; Shaham et al., 2003). Although we did not find strain effects in the seven founder strains that advanced to this final stage of the IVSA protocol, in the larger sampling of CC strains, we found that strains differed in the magnitude of cued reinstatement. Stronger strain effects have been observed during re-acquisition involving reintroduction of both the cue and the drug (Roberts et al., 2018), yet the response to the reintroduction to cues in absence of the drug models a distinct aspect of SUD. The strain differences we observed in the cued reinstatement phase could have multiple sources, including variation in levels of Pavlovian conditioning of the drug-paired (Dickson, McNaughton, et al., 2015) and/or differences in the incentive motivational properties in the reward-associated cue (Bailey et al., 2023). Finally, we found that extinction and cued reinstatement traits are not correlated with more commonly assayed measures of acquisition or cocaine intake, highlighting the importance of modeling these later stages in IVSA studies, yet the striking positive correlation between dose response and cued reinstatement indicates that there are individual differences and genetic variation in the persistence of acquired drug seeking behavior when the effect of the drug is absent, minimal or aversive.

Two important clinical features of SUD are continued drug use despite adverse consequences, and a preference for drug over food and other rewards. These aspects of compulsive drug seeking are poorly captured even by a thorough IVSA protocol (de Guglielmo et al., 2024). Confirmatory testing of high and low drug seeking strains reveal differences in the extent to which they self-administer cocaine when offered with an alternative for food reward, or in the presence of aversive stimuli. The demonstration of these responses in mice suggests the utility of more extensive genetic profiling using these assays to identify strains that might serve as strong preclinical models for compulsive drug seeking.

Human genetics is slowly uncovering genetic variants associated with CUD, typically in people with long and varied history of cocaine use, but the specific role of these variants in particular aspects of cocaine use is challenging to discern from human studies alone. As a result, it is difficult to use this information to inform therapeutic interventions and preventive measures without orthogonal evidence including that from model organisms. Across multiple stages of the IVSA paradigm, we found that traits pertaining to distinct stages, including dose-response, extinction, and reinstatement were variably correlated across the CC strains, and similarly, district traits within the acquisition stage are also highly correlated with one another. Among the founders, PWK/PhJ achieved acquisition criteria in the fewest number of sessions (< 10 sessions) and showed the highest AUC, whereas WSB/EiJ required more than 20 sessions to acquire stable self-administration, and showed the lowest AUC, highlighting a relationship between acquisition and dose response, and indicating that PWK/PhJ is a more vulnerable model than WSB/EiJ to the reinforcing effects of cocaine. The observed relationships among distinct drug-seeking traits obtained across the IVSA paradigm suggests that subsets of these traits may share some of the same underlying processes, whereas others are associated with separate regulatory mechanisms. As we and others have recently shown, functionally relevant genes and pathways discovered in mouse can be used to refine the characterization of human genetic variants associated with CUD through a variety of approaches (Ball et al., 2024; Benca-Bachman et al., 2023; Huggett et al., 2021; Ikeda et al., 2022; Johnson et al., 2024; Reynolds et al., 2021).

Mouse genetics and genetic reference populations, including CC stains and the J:DO population, provide a powerful means of discovering and characterizing neurobiological mechanisms that are shared across multiple risk factors and patterns of drug use. Using advanced, high diversity mouse populations in our IVSA paradigm, we observed a wide range of behaviors that model CUD traits across strains, indicating a genetic basis for the phenotypic differences observed across different stages of CUD. Much has been learned about the mechanisms of cocaine response from C57BL6/J strains and their derivatives, including many of the widely used strains that drive modern neuroscience. However, our findings highlight that the CC population harbors strains that are compelling preclinical pharmacology research models that recapitulate key features of substance use disorders in more substantially than C57BL/6J. Efforts are being made to extend genetic engineering and other capabilities to a broader collection of background strains, and these will allow new research tools and models to be developed (Low et al., 2022). The CC population allows correlation of IVSA related measures with all other behavioral and biological characteristics in this panel, enabling a more complete characterization of these model strains, and for detection of the shared mechanisms of drug self-administration and predisposing behaviors, possibly indicating opportunities for using simpler preclinical assays of cocaine response and susceptibility to SUD. The high precision J:DO population enables the precise mapping of causal variants underlying these behaviors (Logan et al., 2013; Saul et al., 2019). The tremendous variation in shared and disparate patterns of drug self-administration traits across mouse strains provides evidence that these populations will be useful in the identification of specific sources of susceptibility to SUD. Because different strains exhibit different and dissociable patterns of vulnerability as manifest in distinct facets of drug self-administration, specific subtypes of behavior and their effects on the trajectory of SUD-related behaviors can be examined, modeled, and genetically dissected in high-diversity mouse populations. The extreme variation, segregating in the genetically well-randomized population provides an unbiased means of detection of the molecular basis of variation in the underlying processes of SUD-related behaviors.

## Acknowledgements

The authors gratefully acknowledge support from NIDA R01 DA037927 and NIDA P50 DA039841 to EJC. PED was funded by NIDA K99 DA043573 during data collection and early preparation of this manuscript. Cocaine hydrochloride was provided by the NIDA Drug Supply Program. The authors gratefully acknowledge Dr. Stephen Krasinski for assistance with manuscript preparation, and Surgical Services at The Jackson Laboratory for technical assistance with jugular catheter implantation. Dr. Stacey Rizzo for standardizing experimental protocols and training. Dr Marco Venniro and Yavin Shaham consulted on the Choice and Punishment assays. Dr. Vivek Philip, Matt Dunn and others in the Computational Sciences service are partially supported by Cancer Center Support Grant NIH/NCI P30 CA034196. The Mouse Phenome Database, the repository for these data is supported by Dr. Molly Bogue NIDA DA028420.

## Authors contribution statement

EC and PD designed the IVSA experiment. UD designed the choice and punishment assays. PD developed the intravenous cocaine self-administration pipeline in EC’s lab with technical assistance from TR and TW. Mice were tested by TR, TW, ML, AA. LG and JB coordinated mouse colonies and testing. JJ, JB and LT provided technical, analysis and interpretation support related to IVSA studies. MD provided technical assistance with software engineering and data transformation. TW, RB, HF, AK and MS performed statistical analyses in consultation with VP and EC. TW created figures. UD and PD drafted the manuscript which was edited by EC and HF. All authors approved the final version of the manuscript.

