## Supplemental Figures for "Behavioral variation across multiple phases of intravenous cocaine self-administration among genetically diverse mouse populations"

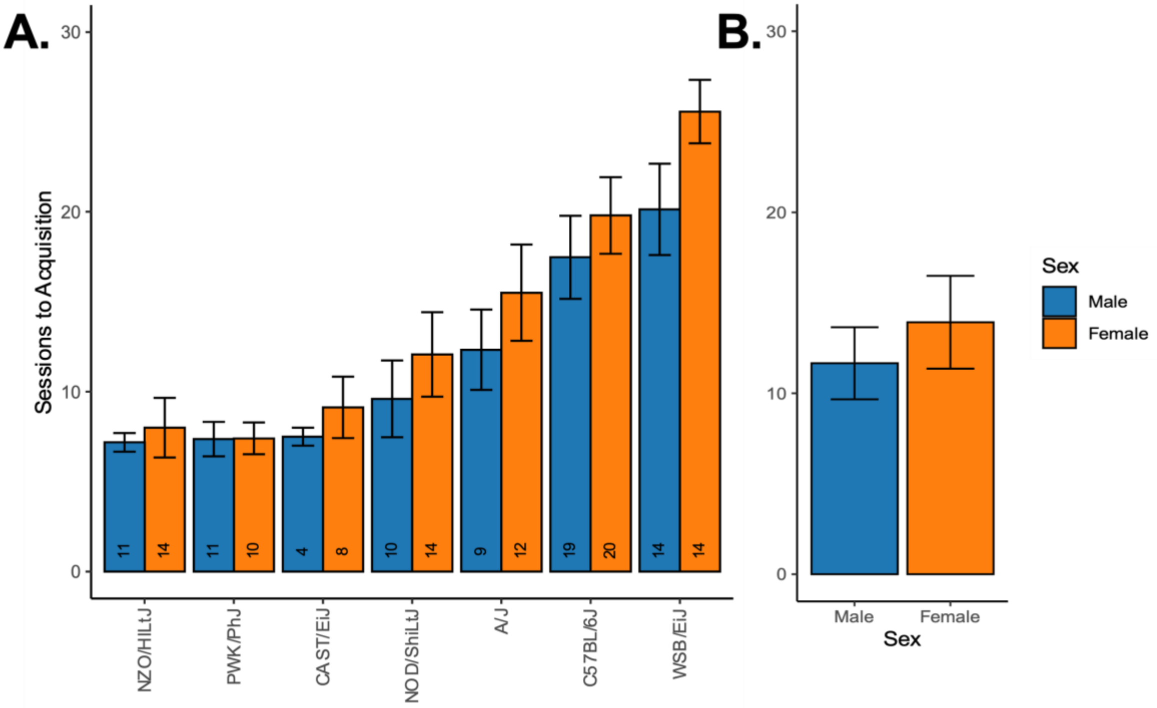


Figure S1. Sex differences in cocaine IVSA acquisition within founder strains. Mean ± standard error (SE) of the number of sessions until acquisition criteria were met in (A) founder strains and (B) across all strains. Note, if acquisition criteria were never met, the maximum number of sessions that mice were exposed to the IVSA paradigm (28 sessions) are reported. Females are denoted in orange and males in blue color. The number of mice per founder strain are listed within each bar.

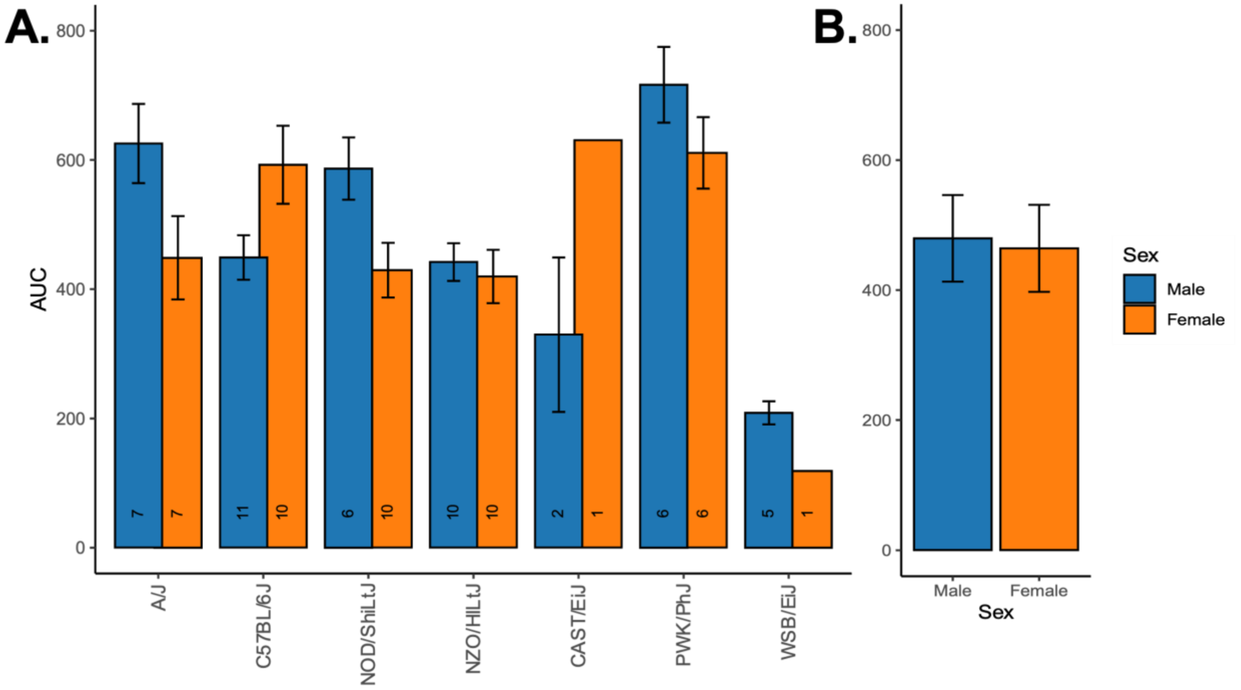


Figure S2. Sex differences in cocaine IVSA dose response across founder strains. Mean ± SE area under the curve (AUC) calculated across multiple dose categories in (A) founder strains and (B) across all strains. Females are denoted in orange and males in blue color. The number of mice per founder strain are listed within each bar.

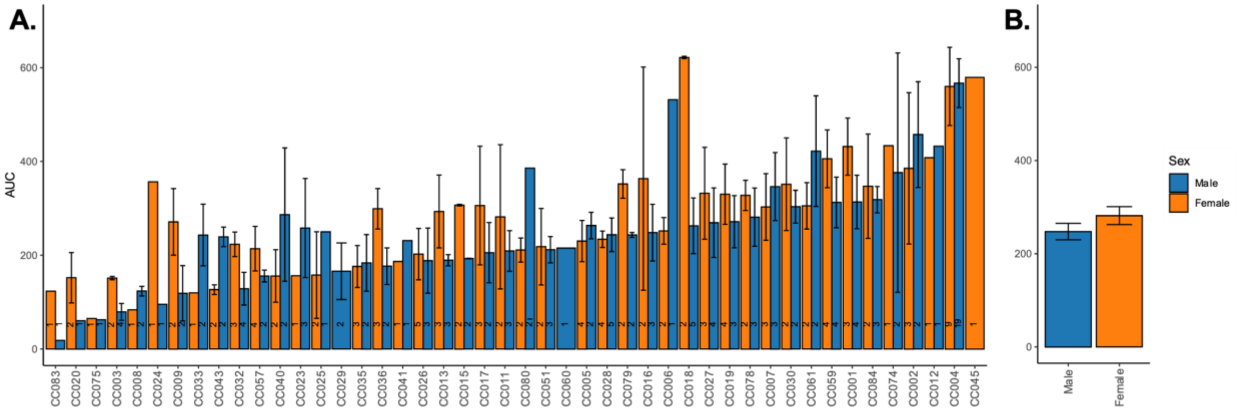


Figure S3. Sex differences in cocaine IVSA dose response across all CC strains. Mean ± SE area under the curve (AUC) calculated across multiple dose categories in (A) CC strains and (B) across all strains. Females are denoted in orange and males in blue color. The number of mice per founder strain are listed within each bar.

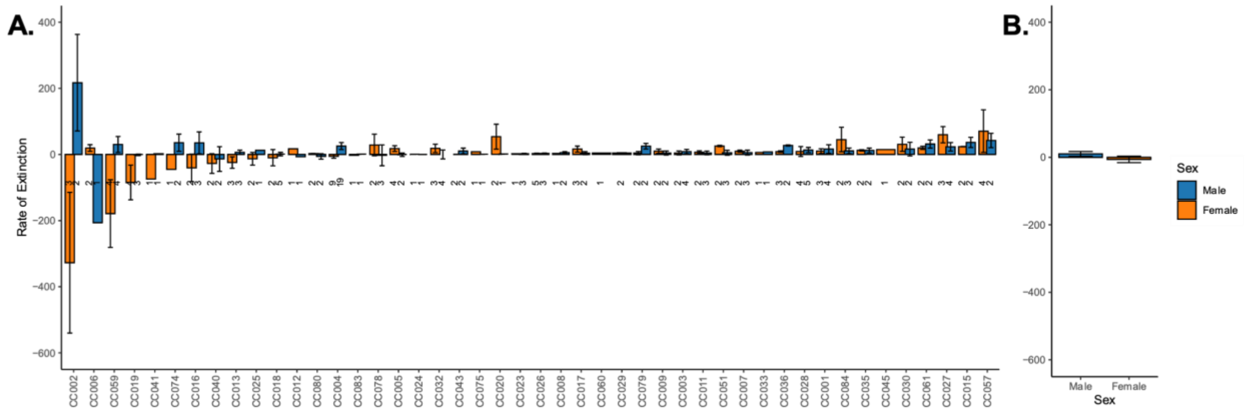


Figure S4. Sex differences in cocaine IVSA extinction across CC strains. Mean ± SE rate of extinction representing change in active lever pressing across extinction sessions in (A) CC strains and (B) across all strains. Females are denoted in orange and males in blue color. The number of mice per founder strain are listed within each bar.
